# Light stimuli and circadian clock affect neural development in *Drosophila melanogaster*

**DOI:** 10.1101/2020.08.07.241208

**Authors:** Eleni Dapergola, Pamela Menegazzi, Thomas Raabe, Anna Hovhanyan

## Abstract

Endogenous clocks enable organisms to adapt their physiology and behavior to daily variation in environmental conditions. Metabolic processes in cyanobacteria to humans are effected by the circadian clock, and its dysregulation causes metabolic disorders. In mouse and *Drosophila* were shown that the circadian clock directs translation of factors involved in ribosome biogenesis and synchronizes protein synthesis. However, the role of clocks in *Drosophila* neurogenesis and the potential impact of clock impairment on neural circuit formation and function is less understood. Here we demonstrate that light stimuli or circadian clock causes a defect in neural stem cell growth and proliferation accompanied by reduced nucleolar size. Further, we define that light and clock independently affect the InR/TOR growth regulatory pathway due to the effect on regulators of protein biosynthesis. Altogether, these data suggest that alterations in growth regulatory pathways induced by light and clock are associated with impaired neural development.

## Introduction

Endogenous circadian clocks are highly conserved and enable organisms to adjust their physiology and behaviour to the day/night cycle. All circadian clocks 1) synchronize to the environment through input pathways, 2) rely on central molecular oscillators, which generate the rhythm and thereby keep circadian time, and 3) transmit time information to modulate the organism behaviour and physiology through output pathways. Processes modulated by the circadian clock include feeding behaviour, locomotor activity, body temperature, hormone level, and metabolic activity (Allada & Chung 2010, Dubowy & Sehgal 2017, Green et al. 2008).

TheA hierarchical network of clocks located in different tissues controls all these rhythmic processes. The master clock is located in the central nervous system (CNS) and synchronizes organ and tissue clocks (Glossop & Hardin 2002, Green et al. 2008). The mammalian master clock is in the suprachiasmatic nuclei of the hypothalamus and consists of ≈ 15000 clock neurons (Liu et al. 2007), whereas in *Drosophila* the master clock consists of a group of 150 clock neurons located in the lateral and dorsal brain (Hermann-Luibl & Helfrich-Förster 2015). The molecular clock machinery is conserved across different species and consists of transcriptional-translational feedback loops (TTFL) to maintain the rhythmic cycling of gene expression (Patke et al. 2020). Briefly, circadian activators trigger transcription of repressor genes which, upon translation, feedback to suppress their own transcription. In *Drosophila*, these circadian activators are Clock (CLK) and Cycle (CYC), which form a heterodimeric protein complex to trigger the transcription of the circadian repressors, *period* (*per*) and *timeless* (*tim*), as well as many other target genes (Dubowy & Sehgal 2017). PER and TIM accumulate in the cytoplasm (Allada & Chung 2010, Dubowy & Sehgal 2017). A complex interplay of light-dependent degradation, initiated by the blue light photoreceptor CRYPTOCHROME (CRY) (Collins et al. 2006, Emery et al. 1998, Emery et al. 2000), and multiple phosphorylation events regulate the accumulation of PER/TIM heterodimers and their timely translocation into the nucleus to inhibit the transcriptional activity of CLK/CYC (Hardin & Panda 2013). Most core clock components are transcriptional regulators modulating the expression of many clock controlled genes in a tissue specific manner (Abruzzi et al. 2011, Rey. et al. 2011, Storch et al. 2002, McCarthy et al. 2007).

The clock in the CNS regulates different biological rhythms of fly, e.g. locomotor activity, sleep, eclosion, whereas circadian control of metabolism mostly depends on peripheral oscillators (Allada & Chung 2010, Green et al. 2008). One of the major peripheral clock in mammals is placed in the liver, which regulates among others metabolism by combining environmental and central clock impulses (Lamia et al. 2008). However, limited food conditions might set the peripheral clock without any involvement of the master clock, indicating that limited nutrition is an external cue to entrain the peripheral oscillators (Damiola et al. 2000, Hara et al. 2001). In *Drosophila*, the fat body takes over the liver function where the clock regulates feeding behaviour and nutrient storage (Xu et al. 2008). In response to nutritional cue, the larval fat body generates mitogen that resumes the cell growth and cell-cycle re-entry in CNS (Britton & Edgar 1998). Thus, regulation of metabolism guaranteeing supply of dietary nutrition plays a key role in the development of the CNS (Sousa-Nunes et al. 2011).

During development, the *Drosophila* CNS is generated from neuroblasts (NB) (Homem & Knoblich 2012, Hakes & Brand 2019), the progenitor cells of the nervous system. Two waves of neurogenesis, embryonic and larval, ensure CNS development and both correlate with changes in NB size. During the embryonic phase NBs go through a limited number of divisions, diminishing in size with each division until they become quiescent. The second wave of neurogenesis, during larval and pupal stages, starts with the exit of NBs from quiescence in a nutrition-dependent manner (Britton & Edgar 1998, Chell & Brand 2010). In contrast to the embryonic NBs, the larval NBs maintain their original size by regrowing after each division. This continues until the end of neurogenesis, where again NBs decrease in size and exit from the cell cycle.

The *Drosophila* central brain has three different types of NBs, which differ by their division modes: Type 0, Type I and Type II. Typically, NBs, like all other stem cells, divide asymmetrically to self-renew and to generate a daughter cell. Daughter cells generated by Type 0 NBs directly differentiate into neurons (Hakes & Brand 2019, Ramon-Cañellas et al. 2018). Contrarily, the division of Type I NBs give rise to ganglion mother cells (GMCs), which divide once more to generate two neurons. Within approximately 100 Type I NBs the four mushroom body (MB) NBs are exceptional, because they proliferate throughout development without a quiescence phase to give rise to the MBs, important structures for the learning and memory processes (Ito & Hotta 1992, Heisenberg 2003, Lin et al. 2013). Type II NBs deposit 12 neurons by each division through intermediate neural progenitors (INPs), which are generated after each division of Type II NBs (Bello et al. 2008, Boone & Doe 2008, Wang et al. 2014, Bowman et al. 2008).

NBs exit quiescence and reactivate proliferation depending on the nutritional state of the organism. The major growth regulator pathway is governed by the insulin receptor (InsR)/ target of rapamycin (TOR) signaling. The pathway is triggered by insulin-like peptide (ILPs) generated in insulin-producing glial cells (IPCs), which receive nutritional signals from the fat body (FB) (Sousa-Nunes et al. 2011, Chell & Brand 2010, Géminard et al. 2009). The TOR pathway might be activated independently from InR pathway, via direct cellular nutrient sensing (González & Hall 2011). The interplay between InsR and TOR pathways regulates cell growth through a variety of effector proteins at the levels of gene expression, ribosome biogenesis, and protein synthesis (Hietakanges & Cohen 2009, Russel et al. 2011).

Approximately 10% of all genes are regulated in a circadian manner (Bass & Takahashi 2010, Doherty and Kay 2010, Wijnen et al. 2006, Ueda et al. 2002, Hughes et al. 2012). Therefore, dysregulation of the circadian system contributes to the pathophysiology of many diseases; most prominent of those are psychiatric disorders and metabolic diseases (Bellet & Sassone-Corsi 2010, Zordan & Sandrelli 2015). Despite the well-established link between the circadian system and physiological processes, the influence of the circadian clock on neuronal development and therefore its potential impact on neural circuit formation is poorly understood. Since the circadian system and cellular signalling pathways are highly conserved from invertebrate to mammals, *Drosophila* is a powerful model for discovering and understanding how developmental processes might be modulated by circadian rhythm. To reveal this link, we compared the neural development of wild-type and clock mutant flies exposed to different light regimes. First, we showed that disturbed circadian clock, as well as different light conditions, affect the NB growth and proliferation. Second, we found that these flies also showed a significant reduction of nucleolus size in the NBs, indicating that rRNA production and further processing might be disturbed, potentially leading to cell proliferation defects. Finally, based on our findings we analysed the effect of light and circadian clock on the InR/TOR growth regulatory pathway and found that both light and clock impair this pathway. Specifically gene expression and activity of Akt, as a regulator of TOR via the InR pathway, and S6K (RPS6-p70-protein kinase) as a downstream target of TOR and major regulator of nutrient metabolism and cell growth, are disturbed when the light regime is changed or clock function is abrogated.

## Results

### Circadian clock and light independently control cell growth in the larval brain

At all developmental stages animals sense and respond to changes in both external and internal conditions, and the combination of this information regulates behaviour and metabolism to benefit from available resources and to maintain cellular homeostasis. Metabolism is important for the proper growth and development of an organism (Koyama et al 2020) and many aspects of metabolism and cellular physiology are controlled by endogenous circadian rhythms (Green et al. 2008, Bellet & Sassone-Corsi 2010, Shi & Zheng 2013). CNS development of *Drosophila* is a prominent example to investigate the mechanism, how an organism copes with changes in environmental conditions, e.g. under nutrient deprivation (Lanet & Maurange 2014). Therefore, we investigated whether disruption of the circadian clock influences the growth of neuroblasts during development. Moreover, we asked whether light, independently from the clock, also affects cell growth. To test the clock effect on cell growth, we measured NBs size in 3^rd^ instar larval brains of wild type and *per*^*01*^ mutants (which lack a functional clock) grown under light-dark (LD) condition. NBs were marked with specific markers (aPKC and Miranda). A similar experiment was done with wild type flies kept in LD, constant darkness (DD, where the molecular clock still maintains a rhythm of approximately 24 hours without being reset by light stimuli) and constant light (LL, where the circadian clock is not functional anymore due to the constant activation of the photopigment CRY). In addition, to control the effects of the light-dark cycles −not directly linked to the resetting of the molecular oscillator by CRY – we included *cry*^*01*^ mutants in our experiments. We found significantly smaller NBs in larvae lacking a functional clock, namely *per*^*01*^ mutants and wild type kept in LL, as well as in larvae with a functional oscillator unable to directly reset by light transitions, i.e. *cry*^*01*^ mutants and wild type flies raised in DD (Figure1B-C). As the strongest effects were observed in *per*^*01*^ mutants and wild type grown under DD condition, all further experiments were performed using these two groups.

In order to distinguish whether the observed growth defect exists from the beginning of the second wave of neurogenesis or it is a later effect, we conducted a staging experiment by measuring the size of NBs during larval development at 22 (1^st^ instar), 36 (2^d^ instar) and 96 (3^rd^ instar) hour after larval hatching (ALH). To analye the late 1^st^ instar larval NBs were chosen, since NB resume reactivation in a couple of hours after hatching and sequentially continue it till the end of 1^st^ instar larval stage. A significant cell growth defect was observed from the late 1^st^ instar larval stage (Figure 1D) much more pronounced at later stages. Thus, our findings indicate that not only the endogenous clock but also light as an environmental factor are needed to control the proper NB growth in the larval brain.

**Figure 1.**
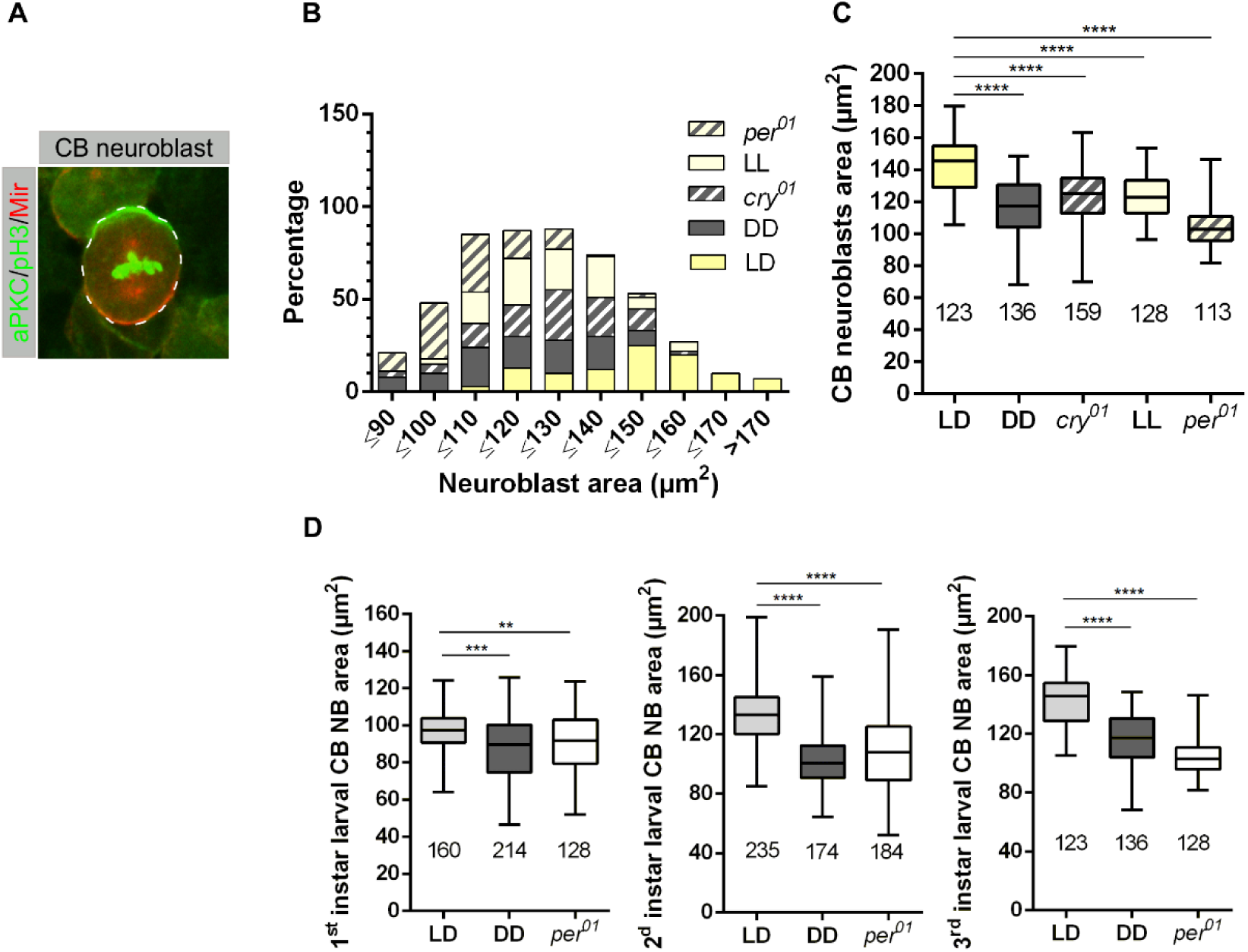
Light and circadian clock control neuroblast growth. **A**) Neuroblasts were marked using aPKC (green) and Mir (red), Phospho-histone H3 (pH3, green) was used to mark chromatin during mitosis; **B**) central brain NBs size distribution of wild-type larvae grown at different light regimes, *cry*^*01*^ and *per*^*01*^ mutant lines (number of measured neuroblasts for each genotype and light conditions are indicated on Fig 1C) where the NBs in wild type mainly distributed within 120 to 170 µm^2^, NBs in wild-type larvae grown under DD condition and *per*^*01*^ mutants range between 90 to 140µm^2^, whereas cell size for larvae under LL condition and *cry*^*01*^ are distributed between 110 to 150 µm^2^; **C**) Average NBs sizes are significantly reduced in *cry*^*01*^ and *per*^*01*^ mutant flies, which were kept under LD condition, as well as in wild type flies developing under different light conditions (p<0.0001); **D**) Difference in neuroblast sizes were observed at all larval stages (p<0.001, p<0.01). At least 10 brains were analysed for each genotype and different light conditions. Each data represents the mean obtained from the number of measured NBs ± the max and min size distribution. The number of measured NBs are indicated below each box.

### Circadian clock and light are required for cell proliferation

The balance between cell growth and proliferation is important to maintain the proper generation, as well as repair of tissues (Maurange et al. 2008). Since we found evidence that the endogenous clock and light have an impact on growth of NBs, we assumed that the proliferation might be affected by the clock and light regime. Therefore, we determined the number of progeny cells for two different subtypes of NBs: mushroom body (MB) and Type II NBs. Since the number of these NBs are only four and eight in each hemisphere, respectively, it is possible to follow the generation of progenies using specific markers. MB and Type II NBs have different modes of division. MB NBs are dividing asymmetrically to give rise to a self-renewing NB and a GMC, which divides one more time to generate two neurons (Figure 2A). Type II NBs divide asymmetrically to generate a self-renewing NB and a transient amplifying cell called immature intermediate neural progenitor (iINP), which by transcriptional changes becomes a mature INP (mINP). Each mINP divides asymmetrically three to five times to form another mINP and a GMC giving rise to two neurons (Figure 2B). To identify MB NBs and their progenies, in addition to Miranda (Mir) as general NB marker, anti-Tailless (Tll) and anti-Dachshund (Dac) were used to specifically label MB lineages (Kraft et al. 2016, Kurusu et al. 2009, Kurusu et al. 2000). Tll is specific for GMCs originating from MB NBs and Dac is a marker for derived neurons (Kenyon cells) surrounding MB NBs. To distinguish Type II NBs, UAS::*mCD8-GFP* was expressed under the control of *wor-*Gal4, *ase*-Gal80 (Neumüller et al. 2011). Additionally, anti-Deadpan (Dpn) and anti-Asense (Ase) antibodies were used to differentiate between progenies of Type II NBs, as Dpn is expressed only in mINPs and Type II NBs, whereas Ase labels iINPs, GMCs and is also co-expressed in mINPs (Bowman et al. 2008, Walsh & Doe 2017). To ensure whether the growth defect is restricted to a certain type of NBs or it is a general effect, we also measured NB size. For Immunohistochemical assay 3^rd^ instar larval brains were used. Cell size analysis revealed that both types of NBs are significantly reduced in size when compared to wild type NBs (Figure 2C and E). This indicates that light input and a functional clock are required for all NBs to maintain proper cell size. For an analysis of the proliferation capacity of MB or Type II NBs, we counted Tll^+^ GMCs and Dpn^+^ INPs generated from single NBs, respectively. As it is shown in Figure 2D and F, in both cases there is a pronounced reduction in the number of progenies. These results demonstrate that the circadian clock and light are required for control of the proliferation of central brain NBs.

**Figure 2.**
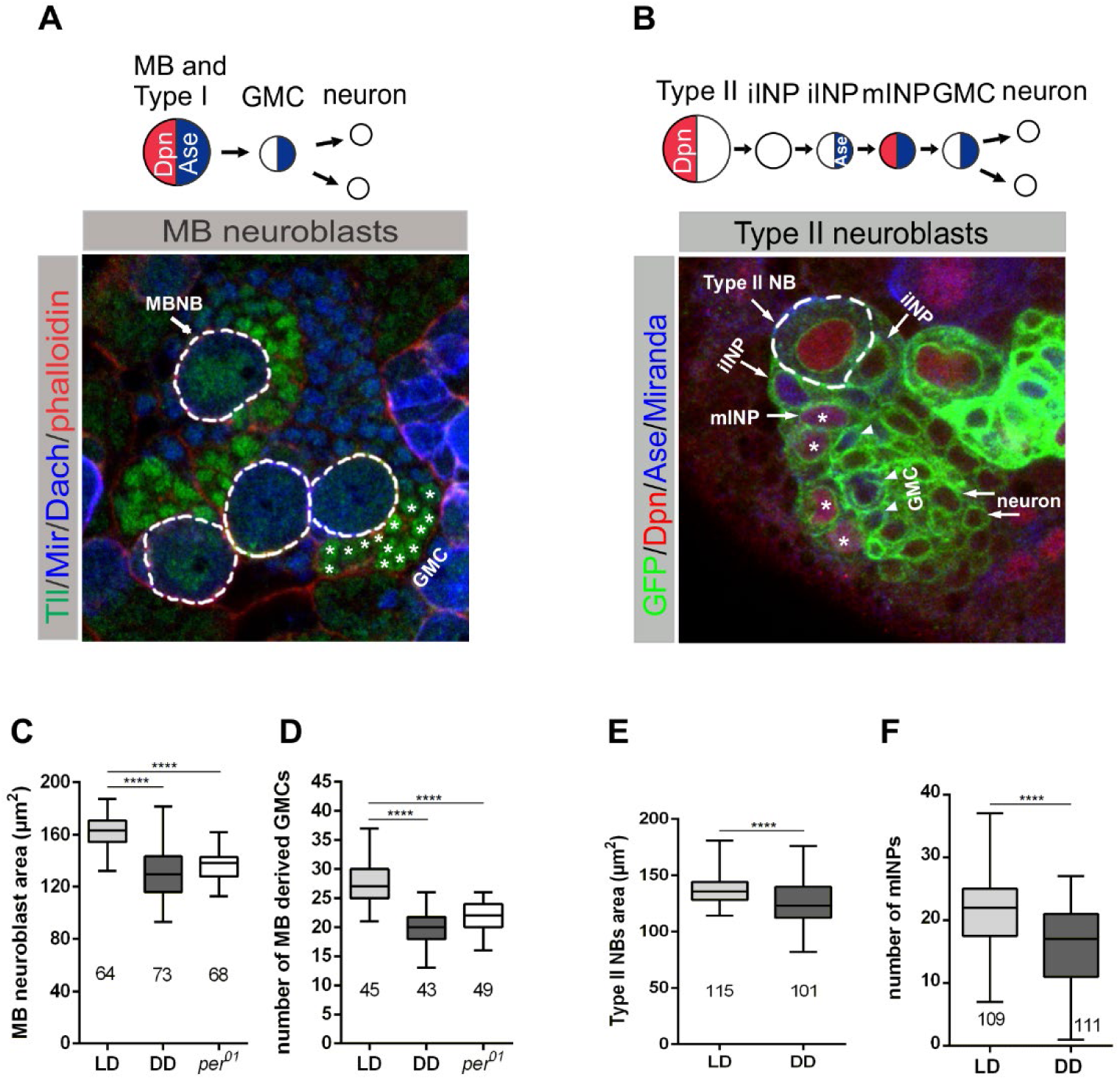
Effect of light and the molecular clock on MB and Type II neuroblast sizes and proliferation pattern. **A**) Miranda (Mir-blue) was used as a neuroblast marker, Tailles (Tll-green) and Dachshund (Dac-blue) were used to mark MB NB derived Kenyon cells and GMCs, correspondingly, and phalloidin (red) was used as a marker for actin. There was a significant reduction in MB NB size **C**) under DD (constant darkness) condition and in *per*^*01*^ mutant flies (p<0,0001); **D**) The cell proliferation was also affected according to the number of Tll^+^ cells (p<0,0001); **B**) Type II NBs were marked with GFP (green) and in addition with NB specific marker Miranda, Dpn (red) and Ase (blue) were used to distinguish progenies of Type II NBs. Type II neuroblast sizes **E**) as well as the number of mINPs **F**) were significantly reduced in flies under DD compare to LD conditions, according to the number of Dpn^+^ cells (p<0,0001). At least 10 brains were analysed for each genotype and different light conditions; each data represents the mean obtained from the number of measured NBs or counted progenies ± the max and min size distribution. The number of measured NBs or counted progenies are indicated below each box.

### Effect of endogenous clock and light on nucleolar size and transcriptional activity

A mechanism, that potentially is responsible for reduced size and consequently proliferation defects involves the disturbance of cell asymmetry and spindle misorientation. Mainly cell-intrinsic processes control NB division. An apical-basal polarity axis is established by the apical enrichment of the Par protein complex components Bazooka, Par6, and aPKC mediating also proper spindle orientation along apico-basal axis. In addition, the Par complex controls basal enrichment of cell fate determinants (the Miranda (Mir)-Brain tumor (Brat)-Prospero (Pros) and Numb-Partner of Numb (Pon) complexes) specifying GMC fate. Upon asymmetric division, apical Par-complex proteins are inherited by the self-renewed NBs whereas basally localied factors are inherited exclusively by the GMC (Homem & Knoblich 2012, Hakes & Brand 2019, Chia et al. 2008). Staining for aPKC and Mir to identify apical and basal cortex (together with phospho-histone H3 (pH3) as a mitotic marker) revealed that central elements of asymmetric cell division (cell polarity and spindle orientation) are not affected (Figure 3A) for each tested genotype and light regime. Hence, impaired cell growth and proliferation is not due to disturbance cell polarity.

**Figure 3.**
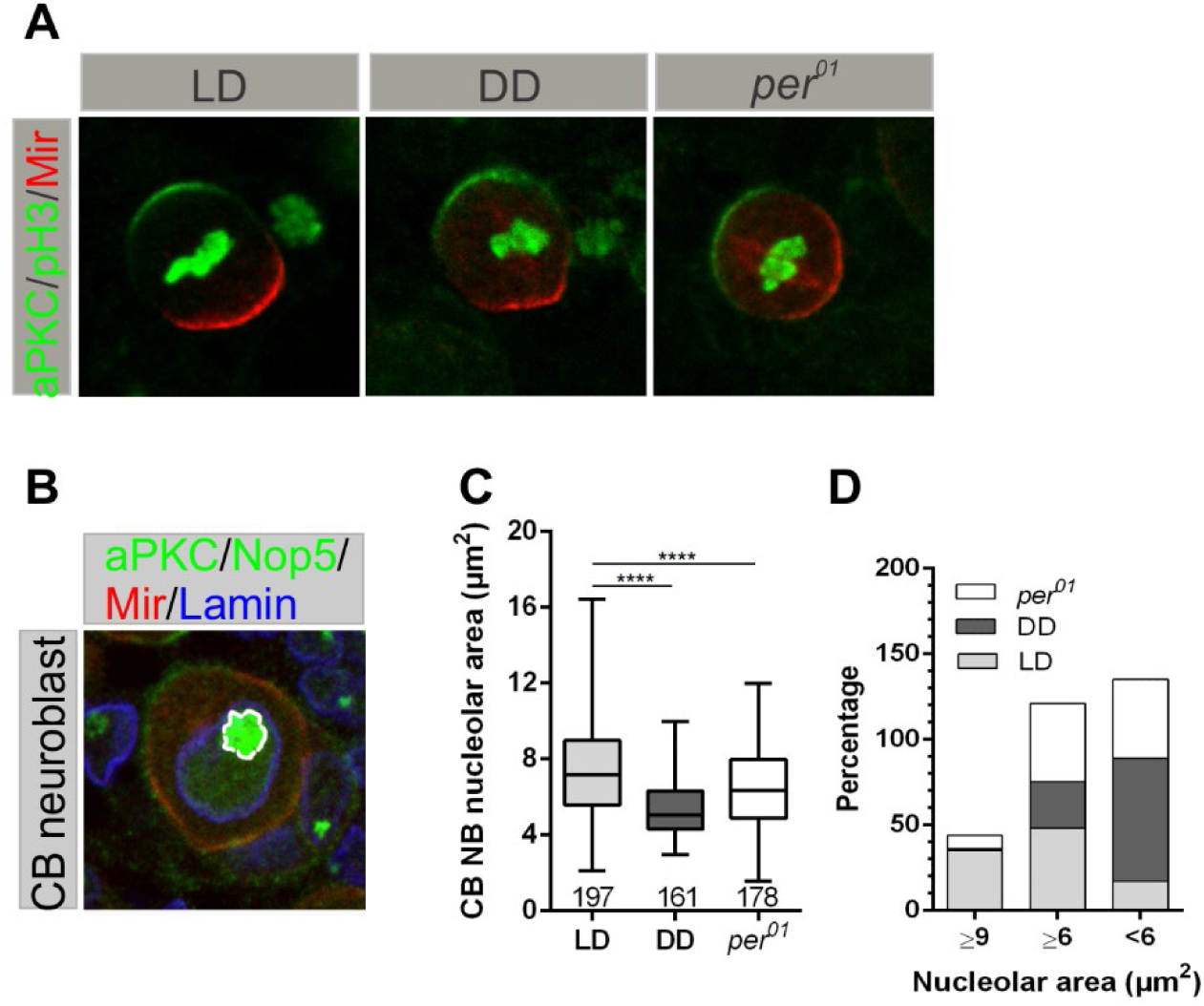
Nucleolar size is affected by light and the circadian clock. **A**) Apical and basal cortical localization of aPKC (green) and Miranda (Mir-red) (as markers for cell polarity) in NB were not disturbed in *per*^*01*^ mutants or wild-type animals grown under DD condition. Phospho-histone H3 (pH3) was used to mark chromatin during mitosis. Correct alignment of chromatin with respect to the apico-basal axis indicates proper spindle orientation upon cell division axis. At least 10 brains were analysed for each genotype and different light conditions; **B**) Neuroblasts were marked using aPKC (green) and Mir (red), Lamine (blue) was used to mark nuclear membranes and Nop5 (green) was used as a nucleolar marker; **C**) ucleolar sizes are significantly reduced in wild-type grown under DD conditions and *per*^*01*^ mutants compare to wild-type in LD situation (p<0.0001). At least 10 brains were analysed for each genotype and different light conditions; each data represents the mean obtained from the number of measured nucleolus ± the max and min size distribution. The number of analysed neuroblasts are indicated below of each box; **D**) Nucleolar size distribution in *per*^*01*^ mutant and wild-type larvae grown under DD and LD light conditions.

Another possible explanation for the compromised proliferation observed in MB and Type II NBs (Figure 2D & F) could be the NBs inability to produce enough proteins to fulfill the requirements of the cell to gain appropriate mass and size before division. The efficiency of protein biosynthesis depends on proper nucleolar function as a site for rRNA transcription/proccesing and assembly of ribosomes. Thus, the nucleolus is a critical player to maintain cell homeostasis and directly affects cell growth and proliferation. Nucleolar size positively correlates with the amount of rRNA biosynthesis (Stepinski 2018). Specifically, it is directly related to RNA polymerase I transcriptional activity and nucleolar size is the largest at the end of G2 phase, before cell division take place (Maszewski & Kwiatkowska 1984, Hernandez-Verdun 2011). To verify, whether light or disturbed circadian clock influence RNA synthesis, we compared the size of NBs nucleoli in *per*^*01*^ mutant and wild type grown under DD and LD conditions. NBs were marked using aPKC (green) and Mir (red), Lamin (blue) to outline the nuclear membrane and Nop5 (green) was used as a nucleolar marker.

To find out whether light or clock affects nucleolar size we analyze interphase cells because of the presence of the nucleolar membrane. Nucleolar size of NBs in *per*^*01*^ mutant and wild-type larvae grown under DD conditions were significantly smaller compared to wild type (Figure 3C). Moreover, classification of nucleolar size, as large (≥9 µm^2^), intermediate (≥6 µm^2^) and small (<6 µm^2^), revealed the strongest effect in wild-type larvae grown under DD conditions, where 72% of the nucleoli were small and only 27% of them at intermediate size. In *per*^*01*^ mutants, the nucleolar size was equally (46%) distributed between intermediate and small sizes, whereas the nucleolar size of wild type in LD conditions were mostly distributed between large (42%) and intermediate (48%) sizes, with only 10% nucleoli being small (Figure 3D).

Taken together, these results provide evidence that the capability of NBs of *per*^*01*^ mutants and wild-type animals grown under DD to synthesize proteins might be impaired, which finally could lead to the observed cell growth defects.

### Expression and activation of InR and Tor signalling are controlled by the light regime and the circadian clock

Reactivation of NBs after dormant state and their regrowth after each division to sustain proper CNS development requires InR/TOR signalling pathway (Sousa-Nunes et al. 2011, Chell & Brand 2010). Impaired function of InR/TOR pathway results in reduced cell size and proliferation as it is required for regulation of dS6K activity, a downstream target of TOR growth pathway, and one of the major regulators of nutrient metabolism and cell growth due to its role in ribosome biogenesis and protein translation (Zhang et al. 2000, Raught et al 2004, Shahbazian et al 2006). Our findings concerning NB and nucleolar size reduction and the role of nucleolus in the maintenance of cell homeostasis lead to the assumption that light might have a regulatory effect on InR/TOR growth pathways. Therefore, we tested the impact of the light on transcriptional and post-transcriptional regulation of dAkt, which is an important signalling molecule in InR pathway (Potter et al. 2002, Ruggero & Sonenberg 2005, Nave et al. 1999). Next, we tested dS6k, a critical effector of TOR growth pathway, which is important for a high level of protein synthesis due to its phosphorylation. First, we checked whether the transcription of *dAkt* and *dS6k* is disturbed under DD condition. Quantitative reverse transcription (RT)-PCR analyses showed that *dAkt* and *dS6k* mRNA transcriptional levels are reduced at most time points during subjective day and night. However, there are no significant differences between flies grown under LD and DD conditions (Figure 4A and C).

**Figure 4.**
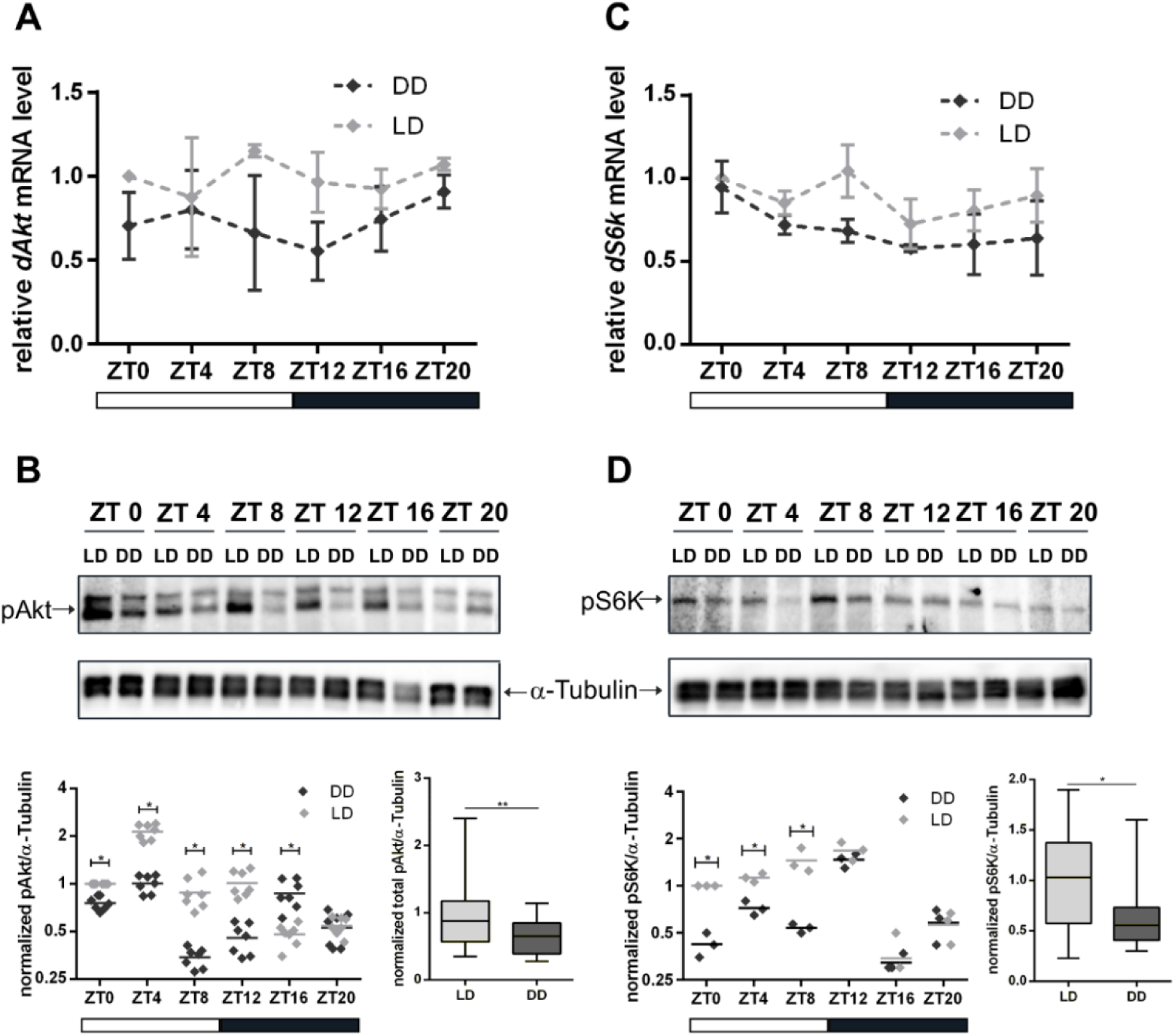
Influence of light on transcription and activation of dAkt and dS6k kinases. **A**) and **C**) expression of *dAkt* and *dS6k* within 24-hour time course under LD and DD conditions. Each time point represents the mean ± standard error of the mean (SEM) obtained from three biological replicates, each repeated in triplicates. As an internal control *rp49* was used; **B**) and **D**) representative blots of 3^d^ instar larval brain extract for dpAkt and dpS6k at different time points during 24-hour, which were grown under different light regimes. α-Tubulin was used as a loading control. Dot plots show measurements of dAkt and dS6k phosphorylation level out of seven and three biological replicates respectively, normalized to α-Tubulin. Box plots display the sum of all time points measurements of dAkt and dS6K phosphorylation for LD and DD conditions (p<0.001 and p<0.01).

In the further test, we did not observe any differences in dAkt protein total expression level during a period of 24 hours between two experimental groups (Figure 4B, Figure 4-figure supplementary 1). Interestingly, although there was no change at transcriptional and total protein expression levels, significant differences were evident in phosphorylation of dAkt and dS6k at most of the time points under DD conditions. Both kinases showed a significant reduction of the total phosphorylation level (Figure 4B and D) resulting in their reduced activity.

We checked also the regulatory effect of the circadian clock on gene and protein expression levels of dAkt and dS6k. Similarly, to the DD condition we did not observe differences in dAkt protein total expression level between wild type and *per*^*01*^ (Figure 5B, Figure 5-figure supplementary 1). However, in contrast to the light effect, the RT-qPCR analysis confirmed that the disruption of the circadian clock results in reduced expression of *dAkt* and *dS6k* mRNAs (Figure 5A and C). In addition, the activity of dAkt and dS6k was severely affected in the circadian clock mutant flies compared to DD condition, exhibiting a robust decrease of the phosphorylation level of both kinases in *per*^*01*^ mutants (Figure 5B and D).

**Figure 5.**
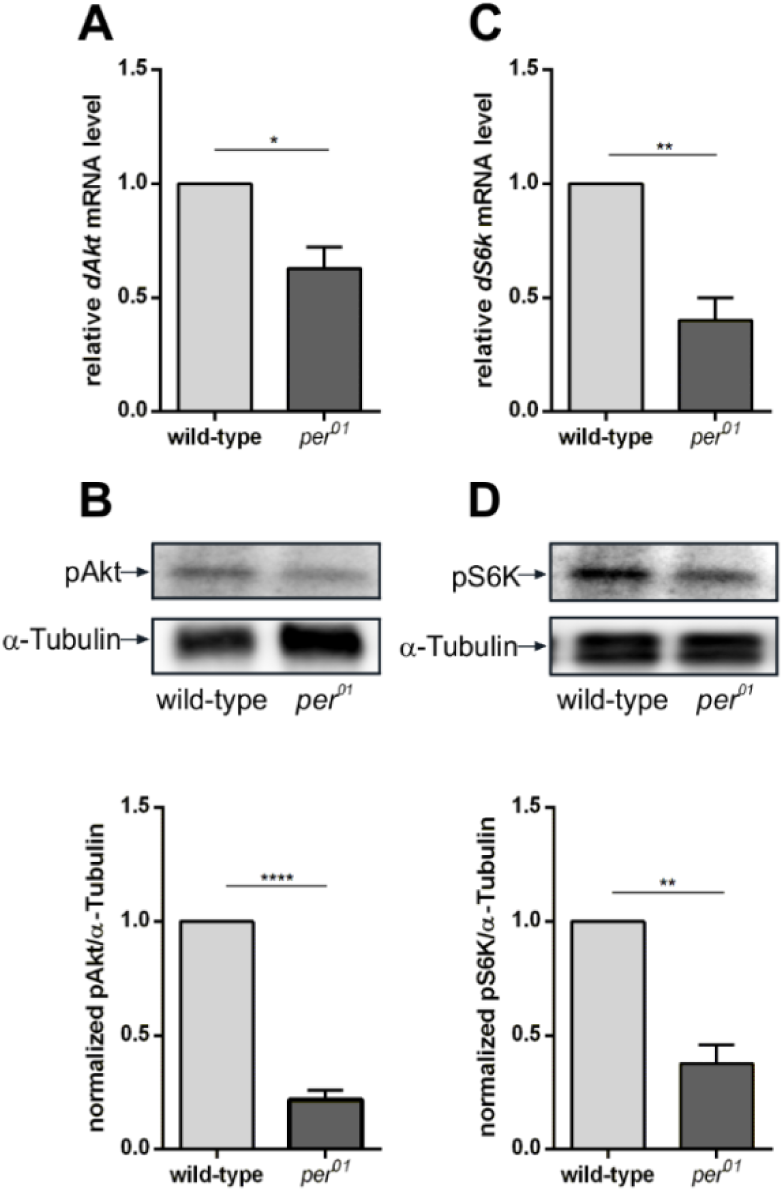
Clock dependent transcription and phosphorylation of dAkt and dS6k kinases. **A**) and **C**) RT-qPCR analysis of *dAkt* (A) and *dS6k* (C) transcription in between wild type and *per*^*01*^ mutant flies. Each bar represents the mean ± standard error of the mean (SEM) obtained from three biological replicates, each repeated in triplicates. As an internal control *rp49* was used (p<0.01 and p<0.004, respectively); Western blots showing dpAkt (**B**) and dpS6k (**D**) phosphorylation levels in wild type and *per*^*01*^ 3^rd^ instar larval brains. α-Tubulin was used as a loading control. Graphs represent measurements of dpAkt and dpS6k phosphorylation level out of three biological replicates normalized to α-Tubulin (mean±SEM) (p<0.0001 and p<0.001).

These results indicate that the activation of two important components of InR/TOR growth pathways is under influence of light and the circadian clock. The impairment of the function of growth pathways via reduction of phosphorylation of dAkt and dS6K might lead to cell growth and proliferation suppression.

## Discussion

### Regulation of neuroblast growth and proliferation by light and the circadian clock

Metabolic homeostasis relies on accurate circadian timing (Green et al 2008, Eckel-Mahan & Sassone-Coesi 2013). The circadian clock is conserved across different species and maintains the rhythmic cycling of gene expression (Patke et al 2020). Many aspects of metabolism and cellular physiology are controlled by endogenous circadian rhythms. In mammals, there is evidence that a disrupted circadian clock causes severe disturbances in rhythmic gene expression (Bellet & Sassone-Corsi, 2010), many of them being involved in metabolism (Panda et al, 2002). Furthermore, environmental factors such as light, temperature or nutrition can externally modulate these autonomous rhythms and synchronize the endogenous clock with the environment. Animals with disrupted circadian system, caused by the exposure to shifted light/dark cycles, show various pathological symptoms including metabolic deficits (Mota et al. 2017, Maury 2019, Barclay et al. 2012, Marcheva et al. 2010) and cognitive difficulties (Gerstner et al. 2008).

Cell proliferation and growth are tightly coordinated to ensure proper animal development. In *Drosophila*, the two waves of neurogenesis (embryonic and postembryonic) are separated by a quiescence state. After exiting from quiescence, NBs first enlarge before they start to proliferate (Truman & Bate 1988, Colombani et *al.* 2003, Chell & Brand 2010). Re-entry to the cell cycle fails when NBs are unable to enlarge due to the lack of nutrition (Britton and Edgar 1998). Nutrient dependent reactivation of NBs is a good model to show how stem cells get adapted to environmental changes. In this study we show that NB growth and proliferation require both clock function and light information. In particular, we demonstrate that different types of CB NBs fail to reach proper size in larvae raised without light (DD) or lack a functional clock (*per*^*01*^) (Figure 1B, Figure 2A and B) and this phenotype is observed from the beginning of the second wave of neurogenesis (Figure 1D). The major part of adult central nervous system is generated during second wave of neurogenesis. To maintain continuous proliferation and to ensure generation of adult nervous system, postembryonic NBs must grow and regain their size between rounds of divisions (Ito & Hotta 1992, Maurange et *al.* 2008, Siegrist et *al.* 2010). Next, we assessed the requirement of light and the circadian clock in the proper proliferation of cells. We observed significant reduction of number of progenies for flies under DD condition and with disrupted clock (Figure 2D and F). Since the process of asymmetric cell division itself was not affected (Figure 3A), decreased NB size could provide an explanation for impaired NB proliferation. Recent findings suggest that NB reactivation requires nutritional input sensed by the fat body, which in turn activates NB ensheating glial cells by fat body derived signal (FDS) to produce Insulin-like peptides (ILP). Reactivation of NBs requires ILP signals and amino acid uptake, which stimulate the InR/TOR signaling pathways to trigger first growth of the neuroblast before they resume proliferation (Sousa-Nunes et al. 2011, Chell & Brand 2010). On the other hand, glial cells also express Activin-like peptides (ALPs) which have a mitogenic effect on NBs and are important for stimulating NB division (Zhu et al. 2008). In this regard, it will be of interest to find out whether glial expression of ILPs and ALPs are affected under DD condition and in clock mutants.

### Coordination of cellular growth regulatory pathways by light and the circadian clock

Prior to proliferation, cells must gain appropriate mass and size which is directly linked to a high demand for proteins. Major steps for protein biosynthesis are the following: RNA synthesis via rDNA transcription, ribosome biogenesis, translation and post-translational modification. In consideration of the high metabolic cost of RNA synthesis and ribosome biogenesis, protein biosynthesis represents one of the main metabolic activities in growing and dividing cells. In mammals, the circadian clock coordinates ribosome biogenesis on transcriptional and translational levels (Jouffe et al. 2013). In particular, authors have shown that circadian clock coordinates mRNA translation via control of the expression and activation of translation initiation factors. The signalling pathways regulating these processes are rhythmically activated in a clock-dependent manner. Interestingly, translated mRNAs are mostly involved in ribosome biogenesis (Jouffe et al. 2013). The nucleolus is the site where RNA synthesis and ribosome biogenesis takes place, thus the size of the nucleolus correlates with cell growth ability. Our observations that flies under DD condition and flies with depleted clock function have drastically reduced nucleoli (Figure 3C & D) indicated that the observed growth and proliferation defect could be due to the failure of protein biosynthesis.

The major growth regulatory pathway, i.e. InR/TOR, is highly conserved between different organisms (Hietekangas & Cohen 2009). Most of the genes regulated by TOR signaling pathway are involved in rDNA transcription, ribosome biogenesis and translation initiation (Guertin et *al.* 2006, Grewal et al. 2007). Furthermore, *Drosophila* larvae deficient for dTOR show a reduced nucleolar size and developmental arrest (Zhang et al. 2000). The observed cell size phenotype raised the question, how the light and circadian clock might regulate cell growth and proliferation? Do they have a role in cell growth regulated by the InR/TOR pathways?

The TOR activation, as the major effector of InR pathway to regulate protein biosynthesis, requires the Akt-phosphorylation event (Gao & Pan 2001, Potter et al. 2002, Inoki et al. 2002). Independent from InR, the TOR pathway also gets activated trough nutritional or amino acid sensing mechanisms (Shim et al. 2013, Colombani et al. 2003, Kim et al. 2008). In both cases, the growth regulation is triggered by phosphorylation of TOR downstream effector proteins. In this report, our focus was on Akt as a major regulator of TOR pathway via InR signaling and one of its major downstream effector protein ribosomal protein kinase p70 (dS6K). dS6K phosphorylates and activates several substrates essential for mRNA translation initiation and translation efficiency (Holz et al. 2005, Dorrelo et al. 2006, Ma et al. 2008). Additionally, activated dS6k plays a role in small ribosome biogenesis via phosphorylation of ribosomal protein S6 (Hietekangas & Cohen 2009, Saxton & Sabatini 2017).

The findings presented here provide correlation that light and the circadian clock control cell growth and proliferation through regulation of InR/TOR pathways. The expression of *dAkt* and *dS6k* was not rhythmic and their level did not differ significantly between wild-type grown under LD and DD within a 24 hours period (Figure 4A and C). However, the activity of dAkt and dS6K was significantly reduced in flies kept under constant darkness at several time points, during a period of 24 hours (Figure 4B and D). We also found, that disrupted clock in larvae has a stronger effect not only on the activity of dAkt and dS6K (Figure 5B and D) but also on their gene expression level (Figure 5A and C). These observations indicate that suppressed NB growth and proliferation might be a result of disturbed TOR activation by Akt, followed by inhibition of translation initiation or small ribosome subunit biogenesis by S6K.

The survival rate and fitness of an organism are regulated by the ability to adapt to environmental changes by adjusting developmental growth, metabolism and behavior. Dependent on different environmental conditions (temperature, light, nutrient availability), animals develop at different rates to adults with different sizes that suit prevailing environmental conditions (Nijhout 2003, Colombani et al. 2003, Shiengelton et al. 2009, Li & Gong 2015). Nutrition dependent hormonal signaling pathways and the endocrine system regulate these processes (Hietakangas & Cohen 2009, Tennessen & Thummel 2011, Boulan et al. 2015). Several experiments in mice models provide evidence for the requirement of circadian rhythms in metabolic homeostasis and energy metabolism (Marcheva et al. 2010, Paschos 2015, Rudic et al. 2004, Yang et al. 2009). Furthermore, the circadian clock regulates rhythmic ribosome biogenesis, on the transcriptional and translational level, through coordination of rhythmic activation of InR/TOR pathways (Jouffe et al. 2013). Our current findings constitute important communication that the major growth regulatory pathways are orchestrated by clock independent and the clock dependent regulation to coordinate proper organism development.

## Material and methods

### Fly Stocks and Genetics

Flies were maintained at 25°C on standard cornmeal food in a 12 h light–dark (LD) cycle. *Canton Special* (*CS*) was the control and genetic background for clock mutant flies: *per*^01^ (Konopka and Benzer, 1971), also was used in light entrainment experiments. *CS* was used also for light entrainment experiments. To mark Type II NBs, the *wor-*Gal4, *ase*-Gal80 driver line was used (Neumueller et al. 2011) to express UAS::*mCD8-GFP*. For RNA isolation or protein extraction, after laying eggs, vials were entrained in a 12 h LD cycle or shifted into constant darkness until the 3^rd^ instar larvae stage. Relative to Zeitgeber time 0 (ZT0) as the time of lights-on during the LD cycle and circadian time 0 (CT0) as the time corresponding to subjective lights-on during free running in DD, larvae were collected in a ZT0-ZT4-ZT8- ZT12-ZT16-ZT20-CT0-CT4-CT8-CT12-CT16-CT20 schedule and dissected. Isolated larval brains were collected either in Trizol (for RNA extraction) or in Laemmli buffer (for protein extraction).

### Larval staging

Immediately after hatching, larvae were collected from apple juice plates at 30 minutes intervals and transferred to standard fly food plates with yeast. *per*^*01*^ mutants, wild-type larvae under LD and DD light regime were kept at 25°C until they reached the desired age. Before preparation, larval stages were determined by means of spiracle morphology. 36 hours old larvae were L2, 48 hours and older larvae were L3.

### Immunohistochemistry (IHC)

For immunostainings, larval brains were dissected in PBS (10 mM Na2HPO4,2 mM KH2PO4, 2.7 mM KCl, 137 mM NaCl) and fixed on ice for 25 min in PLP solution (2% paraformaldehyde, 10 mM NaIO4, 75 mM lysine, 30 mM sodium phosphate buffer, pH 6.8). All washings were done in PBT (PBS plus 0.3% Triton X-100). After blocking in PBT containing 3% normal goat serum for 2 h, brains were incubated overnight with combinations of the following primary antibodies: rabbit anti-protein kinase C (anti-PKC) clone C20 (1:1000; Santa Cruz Biotechnology), rabbit anti-phospho-histone H3 (1:2500; Millipore Upstate), rat anti-Miranda (1:300; Abcam, clone CD#5-7E9BG5AF4), mouse anti-Miranda (1:20; F. Matsuzaki, Kobe, Japan), rabbit anti-Nop5 (1:600; G. Vorbrüggen, Göttingen, Germany), chicken anti-GFP (1:1000; Abcam), rabbit anti-Asense (1:400; J.Diaz-Benjumea, Madrid, Spain), guinea pig anti-Deadpan (1:1000; J. Knoblich, Vienna, Austria), AlexaFluor546-Phalloidin (1:100; Molecular Probes), mouse anti-Dachshund (1:10; clone mAbdac2-3, Developmental Studies Hybridoma Bank, Iowa City, IA, United States), rabbit anti-Tailless (1:600; J. Reinitz, Chicago, Illinois, United States), guinea pig anti-lamin DmO (1:300; G. Krohne, Wuerzburg, Germany). Secondary antibodies were conjugated with Alexa Fluor 488 (Molecular Probes), Cy3, Cy5, or DyLyte488 (Dianova). After extensive washing in PBT, brains were embedded in Vectashield. Confocal images were collected with a Leica SPE microscope. Image processing was done with ImageJ 1.46r software (NIH, Bethesda, MD).

Neuroblast and nucleolar areas were measured by the freehand selection tool of ImageJ 1.46r software (NIH, Bethesda, MA, USA). Neuroblasts were visualised by aPKC or Miranda, nucleoli by Nop5 antibodies.

### Neuroblast proliferation and IHC analysis

Progenies of MB and Type II neuroblasts were marked with specific antibodies. Particularly, MB NBs daughter cells were marked with anti-Tll antibody, which specifically marks MB NB derived GMCs (Kurusu et al. 2009). Type II NBs and their lineages were marked with GFP by using UAS::*mCD8-GFP* line under the control of the *wor-*Gal4, *ase-*Gal80 driver line (Neumueller et al. 2011). To specify between all types of the Type II NB lineages anti-Dpn and anti-Ase antibodies were used (Bowman et al. 2008, Walsh & Doe 2017). MB NB derived GMCs and mINPs were counted with Amira® software using the Landmark selection.

### RNA isolation and RT-qPCR analysis

Entrained larvae, as well as wild type and *per*^*01*^ mutant larvae were collected in PBS and placed on ice. Dissections of larvae were done within 30 min and brains were collected in Trizol. Total RNA was extracted from brains using TRIzol^®^ reagent according to the manufacturer’s instruction (Ambion^®^, Thermo Fisher Scientific, Waltham, MA, United States). First-strand cDNA was synthesized from 2 µg of RNAusing High-Capacity cDNA Reverse Transcription Kits (Applied Biosystems, Thermo Fisher Scientific, Waltham, MA, United States). RT-qPCR was done using PowerUp™SYBR™Green Master Mix (Applied Biosystems, Thermo Fisher Scientific, Waltham, MA, United States) on a StepOnePlus™ (Applied Biosystems. Thermo Fisher Scientific, Waltham, MA, United States) real-time thermal cycler. Reaction mixtures contained 300 nM of oligonucleotides (see Supplementary Table S1). RT-qPCR conditions were 2 min 50°C and 2 min 95°C holding steps, followed by 40 cycles of 15 s 95°C and 1 min 60°C. Results were expressed as fold change in expression of the treated sample in relation to untreated samples and relative to the reference gene *rp49*. Mean ±SEM was calculated at least from triplicate experiments from each of the three independent biological samples per genotype or different light regime.

The following primers were used to amplify the cDNA of target genes:

*Akt* (forward 5’-ACAGATCTAGTGTTGAAAAAAATATACCG-3’ and reverse 5’- ATGTCTCCTTGGTAGCTGAACTGCG-3’), *S6k* (forward 5’- TTCTTAGAGGATACCACATGCTTC-3’ and reverse 5’- TGGTCAAAATTTCAGGTGCCATGTAC-3’), *rp49* (forward 5’- GCCCAAGATCGTGAAGAAGC-3’ and reverse 5’-CGACGCACTCTGTTGTCG-3’).

### Western Blot

Lysates from wild-type or *per*^*01*^ larval brains were sonicated in 2x Laemmli, separated by SDS-PAGE and transferred to PVDF membranes. Blots were incubated overnight at 4°C with the following antibodies: rabbit anti-phospho-Akt (Ser473) (1:1000; clone D9E, Cell Signaling), rabbit anti-Akt (1:1000, Cell Signaling, Denvers, MA, US), rabbit anti-phospho-Drosophila-p70 S6k (1:1000; Cell Signaling, Denvers, MA, US), mouse anti-α-Tubulin (1:5000, clone NDM1A, Merck, Darmstadt, DE). After incubation with HRP-coupled secondary antibodies, signal detection was done with the ECL Plus detection reagents (GE Healthcare Life Science, Buckinghamshire, UK) and a ChemoCam ECL Imager equipped with a 16 bit camera (Intas, Göttingen, DE). Exposure times were adjusted to allow for quantification of signal intensities within the dynamic range of the camera system.

### Data Analysis

Cell and nucleolar size, as well as proliferation analysis, were analysed in the program Statistica *v*. 9. Distributions of variables did not deviate significantly from normality (Kolmogorov-Smirnov test; *P* > 0.2). A one-way analysis of variance (ANOVA) was performed for statistical analysis. Axis lengths and areas were considered as dependent variables, and the strain (wild type versus mutant or LD light regime versus DD) was considered as an independent variable.

To calculate the significance between expression data differences, statistical analyses were performed using Mann–Whitney-*U*- test (Origin Pro9.0.0 b45 software) to determine significant differences between genotypes. For multiple testing within one data set, the level of significance *p* 0.05 was adjusted with the Bonferroni correction factor. Graphs are presented as mean ± the max and min size distribution or ± SEM, asterisks depict the level of statistical significance \*\*\*\**p* and \*\*\**p* ≤ 0.0001, \*\**p* ≤ 0.001 and \**p* ≤ 0.01. Graphs were generated in Prism6.

## Acknowledgments

We would like to thank Dr. Vahan Serobyan for discussion and comments on the manuscript. We thank Kjara Sophia Pilch for assistance in RT-qPCR experiments.

This work was founded by the German Research Foundation (DFG, SFB 1047 “Insect Timing”, Project A6 - T.R.) and the German Excellence Initiative to the Graduate School of Life Sciences, University of Würzburg (GSC106 - A.H.).

