## Supplementary material for "Light stimuli and circadian clock affect neural development in *Drosophila melanogaster*": Figure 4B, Figure 4-figure supplementary 1

**Supplementary figures**

**Figure 4B, Figure 4-figure supplementary 1**

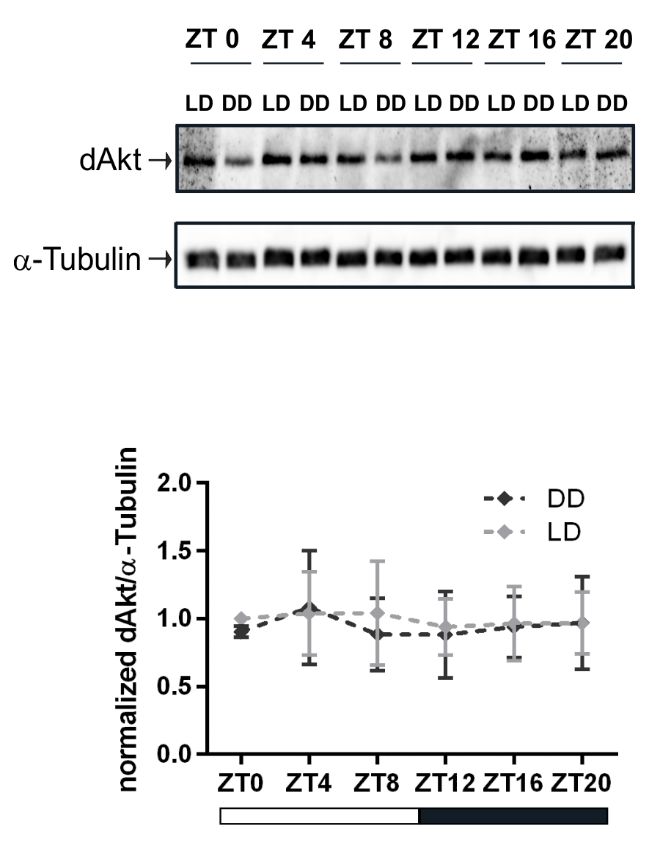

**Figure 4B, Figure 4-figure supplementary 1. Light dependent expression of dAkt.** Representative blot of dAkt expression from wild-type 3^rd^ instar larval brain extract grown under different light regimes at different time points during 24-hour. α-Tubulin was used as a loading control. Graphs show measurements of dAkt expression level out of seven biological replicates, normalized to α-Tubulin (mean±SEM).
