## Supplementary material for "Light stimuli and circadian clock affect neural development in *Drosophila melanogaster*": Figure 5B, Figure 5-figure supplementary 1

**Supplementary figures**

**Figure 5B, Figure 5-figure supplementary 1.**

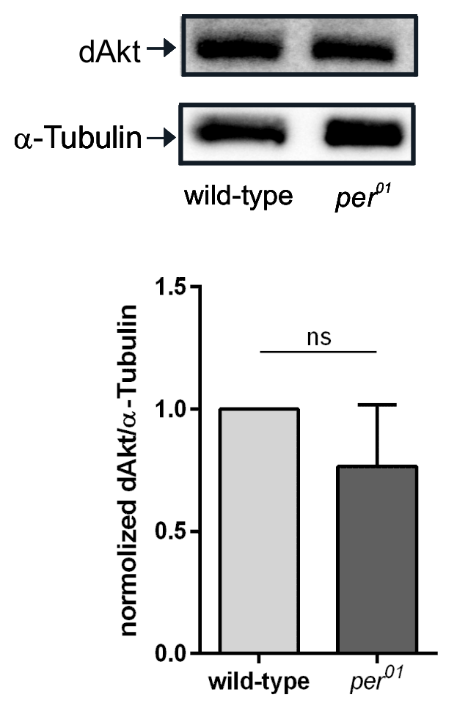

**Figure 5B, Figure 5-figure supplementary 1. Clock dependent expression of dAkt.** Representative blot of dAkt expression from wild-type and *per^01^* mutant 3^rd^ instar larval brain extract at different time points during 24-hour. α-Tubulin was used as a loading control. Graphs show measurements of dAkt expression level out three biological replicates, normalized to α-Tubulin (mean±SEM).
